# Phenotypic and transcriptomic characterization of bicalutamide and enzalutamide resistance in castration-resistant prostate cancer cells

**DOI:** 10.1101/2025.05.30.656961

**Authors:** Reetta Nätkin, Milja I. Paloneva, Paavo V. H. Raittinen, Heimo Syvälä, Merja Bläuer, Teuvo L. J. Tammela, Pauliina Ilmonen, Matti Nykter, Aino Siltari, Teemu J. Murtola

## Abstract

The cornerstone treatment for aggressive prostate cancer (PCa) is androgen deprivation therapy (ADT). Eventually PCa cells develop castration resistance, i.e. resistance to ADT. Castration resistant PCa initially responds to androgen signaling inhibitors such as bicalutamide and enzalutamide, but the cells become resistant to these drugs as well overtime.

We used long-term cell cultures to create testosterone-dependent, testosterone-independent (CT), bicalutamide-resistant (BR), enzalutamide-resistant (ER), and sequential resistant BR-ER VCaP cell lines to investigate transcriptomic and proteomic changes during the development of drug resistance. Phenotypical changes were evaluated based on cellular morphology and sensitivity to docetaxel.

We observed marked changes in transcriptome during the development of drug resistance and differences between ER and sequential BRER. Androgen response and fatty acid metabolism were upregulated in both ER and BRER. MYC target hallmarks were positively enriched only in BRER. Negative regulation of cell death was upregulated in ER while in BRER cell cycle related pathways were upregulated. Direct comparison of ER and BRER revealed androgen signaling signature being higher in ER than in BRER. BRER was more resistant to docetaxel than ER. Clinical significance of our results was confirmed using Finnish patient data and public patient data. In castration resistant patients’ response to enzalutamide treatment were decreased in patients first treated with bicalutamide comparing patients treated only with enzalutamide.

In conclusion, we showed that transcriptomic and phenotypic changes occurring during formation of drug resistances and docetaxel sensitivity depend on the sequence of treatments. Treatment responses were also different in sequentially treated patients. In the future, this may inform treatment sequencing to retain the PCa cells more sensitivity to subsequent treatment lines.

## 1. Introduction

Prostate cancer (PCa) is the second leading cause of cancer-related deaths in men [1]. While prognosis is often favorable, 15-25% of cases progress to metastatic disease [2]. Disease progression is driven by androgens, which activate the androgen receptor (AR) [3]. Androgen deprivation therapy (ADT), reducing testosterone levels to castration levels, is the standard treatment for advanced PCa [4]. However, resistance to ADT eventually develops, leading to castration-resistant prostate cancer (CRPC), with AR signaling reactivation being a key mechanism. AR signaling inhibitors (ARSIs) like bicalutamide and enzalutamide are effective initially, but resistance to these therapies eventually develops [5,6]. Additionally, cytotoxic agents like docetaxel are used for aggressive CRPC.

ADT, ARSIs and docetaxel induce transcriptomic changes in PCa. ADT commonly triggers AR upregulation, while resistance to enzalutamide and bicalutamide is linked to AR upregulation and alternative splicing [7,8]. The transcriptomic alterations driving docetaxel resistance are less well understood [9,10]. It remains unclear whether different AR-targeted therapies create distinct transcriptomic and proteomic profiles post-resistance, or if these changes depend on the sequence of ARSI treatments [11]. Additionally, the impact of these changes on sensitivity to subsequent therapies, like docetaxel, is not fully understood.

We used testosterone-dependent, testosterone-independent, bicalutamide-resistant (BR), enzalutamide-resistant (ER), and sequentially resistant BRER VCaP cell lines, developed in our laboratory. Resistance was induced by continuous treatment until growth resumed. We assessed cell viability under docetaxel treatment and observed morphological changes to evaluate the potential benefits of these adaptations.

## 2. Materials and methods

### 2.1 Materials for cell growth

The VCaP cell line, originally derived from bone metastatic cells of hormone-refractory prostate cancer, was generously provided by Dr. Tapio Visakorpi from the Tampere University in Finland. RPMI 1640, L-glutamine, and antibiotic-antimycotic solution (A/A) were obtained from Invitrogen (Carlsbad, CA, USA). Charcoal-stripped fetal bovine serum (DCC-FBS), testosterone (T), and bicalutamide were purchased from Sigma-Aldrich (St. Louis, MO, USA), while enzalutamide (MDV3100) was obtained from SMS-Gruppen Selleckchem (Rungsted, Denmark). Docetaxel was acquired from Cell Signaling Technology Inc. (Danvers, MA, USA). Cellbind® well plates were obtained from Corning (Corning, NY, USA), and all other disposable cell culture materials were purchased from Nalge Nunc International (Rochester, NY, USA). For SDS-PAGE and subsequent immunoblotting analysis of the indicated proteins, the following reagents were used: Laemmli sample buffer for SDS-PAGE (Sigma), Precision Plus Protein Standards (Bio-Rad Laboratories, Hercules, CA, USA), NuPage transfer buffer (Invitrogen, Carlsbad, CA, USA), horse radish peroxidase-conjugated secondary antibodies (Cell Signaling Technology Inc.), and enhanced chemiluminescence reagents (ECL Western Blotting Detection Reagents, GE Healthcare, Buckinghamshire, UK).

### 2.2 Establishment of subclones of the VCaP cell line

The parental VCaP (RRID: CVCL_2235) cells were cultured in RPMI 1640 supplemented with 10% DCC-FBS, 1% L-glutamine, 1% A/A, and 10 nM T for seven months to establish a T-dependent subclone, VCaP-T cell line. The VCaP-T cells were dependent on the presence of 10 nM T. Subsequently, VCaP-T cells were cultured in the presence of 0.1 nM T for 10 months to establish the subclone VCaP-CT that was able to grow with castrate level of T. VCaP-CT cells were then cultured in the presence of androgen signaling inhibitors (bicalutamide or enzalutamide at concentrations of 10 µM each) for 12 months to establish subclones resistant to the indicated AR signaling inhibitors (subclones VCaP-BR and VCaP-ER, respectively). Moreover, VCaP-BR cells were also treated with enzalutamide at a concentration of 10 µM for 12 months to obtain a subclone that had developed sequential resistance first to bicalutamide and then to enzalutamide (subclone VCaP-BRER). The growth of the cell lines was measured using a modified crystal violet assay, as described in Murtola et al. 2009 [12]. All cell lines used in this study was genotyped using single nucleotide polymorphism -based assay including eight SNPs at March, 2025 and as a result all cell lines had original genotype of VCaP cells.

### 2.3 Testosterone sensitivity analysis

To test T sensitivity, the established cell lines were seeded in 6-well plates with a density of 500,000 cells per well in their respective growth media and allowed to attach for 48 h before the treatments. VCaP-T and VCaP-CT cells were then cultured under increasing concentrations of T (0-100 nM) for seven days before crystal violet staining [12]. VCaP-T cells cultured under 10 nM T were used as a reference when VCaP-T and VCaP-CT cells responses to dividing T concentrations were evaluated. In drug-resistant VCaP-BR, VCaP-ER, and VCaP-BRER cells cultured with 10 µM bicalutamide/enzalutamide and 0.1 nM T were used as controls, and the relative cell growth of other groups was calculated by comparing them to that of the controls. The mean and standard deviation of three separately cultured wells were calculated and visualized using GraphPad Prism (version 9).

### 2.4 SDS-PAGE and Western blot analysis

Protein extraction from each established subclone of parental VCaP cells was performed using M-PER® reagent (Pierce, Rockford, IL) modified with protease inhibitors (Complete Mini Protease Inhibitor Cocktail tablets; Roche Diagnostics GmbH, IN, USA). Total extracted protein concentrations were measured using the BCA Protein Assay Kit (Pierce) according to the manufacturer’s instructions. SDS-PAGE and immunoblotting analysis were performed as described by Murtola et al. 2012 [13]. Immunoblotting was performed by incubating membranes (Trans-Blot Turbo, BioRad) with primary antibodies (AR, 1:2000, ab13373, Abcam; AGR2, 1:200, sc-101211, Santa Cruz; IGF1R, 1:400, NBP1-77679, Novusbio; ITM2A, 1:500, A62806, Biosite; PDCD6, 1:1000, 12303-1-AP, Biosite; TMEFF2, 1:100, MBS668238, SMS-gruppen; NDRG1, 1:200, sc-398291, Santa Cruz; YWHAZ, MBS668238, MyBioSource; Cyclin D, 1:10000, ab134175, Abcam; Calnexin, 1:20000, sc-29954, Santa Cruz; Nkx3.1, 1:100, sc-393190, Santa Cruz; and HSP90, 1:5000, ab203085, Abcam) at 4°C overnight, followed by 1-hour incubation with a secondary antibody (anti-mouse or -rabbit horseradish peroxidase (HRP)-conjugated secondary antibody, Cell Signaling) at room temperature. The densitometric analysis of the immunoreactive protein bands was performed using the ImageLab software (BioRad, Hercules, CA, USA). β-actin (1:3000) or β-tubulin 1 (1:5000, both from Sigma-Aldrich) was used as a loading control. Experiments were repeated three times with protein samples from independent cultures of each indicated cell line. Differences in the reported protein expression between cell lines were calculated as average fold changes using the protein expression levels in the VCaP-CT cell line as a control and visualized using GraphPad Prism (version 9).

### 2.5 RNA-seq and quality assurance

RNA was extracted using Direct-Zol Miniprep Plus Kit (Zymo Research #R2070) according to the manufacturer’s instructions. Library preparation was done according to Illumina TruSeq© Stranded mRNA Sample Preparation Guide (part # 15031047). All cell lines, three replicates of each, were sequenced using Illumina HiSeq 3000 with 75 bp-long paired-end reads. Quality control of the raw read data was performed using FastQC version 0·11·7. Furthermore, the RNA-seq data was aligned with STAR version 2·5·4b [14], using Ensembl reference genome GRCh38. The gene-wise reads were quantified with featureCounts version 1·6·2 [15], and for the transcript variant counts kallisto version 0·44·0 was used [16], along with Gencode annotations release 28.

### 2.6 Cell imaging

Microscopic analysis of the cell morphology and growth characteristics of the cell lines was performed using a Nikon Ti-S inverted phase contrast microscope. For imaging, 500,000 cells of each line were seeded in T25 culture bottles in the presence of their respective T level and AR signaling inhibitor supplements and cultured for six days.

### 2.7 Docetaxel cytotoxicity assay

For docetaxel cytotoxicity studies, the established cell lines were seeded in 6-well plates (500,000 cells per well) in their respective growth media and allowed to attach for 48 h before treatments. The cells were then treated with vehicle alone (dimethyl sulfoxide, DMSO) or with different concentrations of docetaxel (1-10 nM) for 72 h. After incubation, relative cell numbers were assessed using a modified crystal violet method [12]. Growth analyses were performed in six independent cell culture samples of each cell line, and the results are shown as the average growth using the VCaP-CT cell line as a control.

### 2.8. Clinical cohort collection

This retrospective cohort study included 493 Finnish men with castration-resistant prostate cancer who had used second generation antiandrogens (ARSI) i.e. enzalutamide, apalutamide or darolutamide between January 1, 2012, and December 31, 2024. The collected data included also information of previous use and duration of use of bicalutamide and ADT, clinical tumor characteristics, physical performance classifications, and additional comorbidities and medications. In the analysis of response to ARSI treatment after formation of castration resistance, follow-up started at ARSI initiation and ended at death or December 31, 2024. In the analysis of overall response to advanced PCa treatment, follow-up started at the initiation of ADT and ended at death or December 31, 2024. Patient data were collected from the Wellbeing Services County of Pirkanmaa database with research permit R24223.

### 2.9 Statistical analysis

For the analysis of differentially expressed genes (DEGs), the quality-controlled mRNA counts were normalized using the median-of-ratio method with DESeq2 version 1·26·0 in R version 3·6·3 [17]. The statistical significance of DEGs between cell lines was modelled using Negative Binomial regression and Wald test for significance, using a significance level of 0·0001, also implemented with DESeq2 [17]. The p-values were adjusted for multiple comparison errors using the Benjamini-Hochberg method. The analysis was further limited to genes showing a log2 difference greater than 1 and mean of normalized counts greater than 4000 counts across all samples.

The transcriptomic profiles of the genes that passed these threshold values were visualized as Venn-diagrams and a heatmap. In the heatmap the relatively most expressed gene mRNA/cell population is colored as bright orange, whereas the relatively less expressed gene mRNA/cell population is colored as dark purple. For each cell line, the midpoint color was defined as the median of log2-transformed gene expression values.

Gene set enrichment analysis (GSEA) for DEGs was performed with fgsea version 1·10·1 R-package. The hallmark gene set, GO and KEGG version 7·5 gene set annotations were downloaded from MSigDB. Adjusted p-value < 0·05 was considered as significant. For DEG sections specific and common to VCaP-ER and VCaP-BRER, over-representation analysis (ORA) was performed in ConsensusPahtDB release 35 [18]. KEGG pathways and GO categories, excluding level 2 and cellular component categories, were considered. P-value cutoff was set at 0·01. Selected (representative terms among terms fulfilling the following criteria: q-value <= 0·1, generatio > 0·01, and count >= 3) enriched KEGG pathways and GO terms were visualized with ggplot2 version 3·4·2 R-package.

The splice variant distributions were visualized by first normalizing the mRNA counts using the median-ratio method. For variant-variant comparison, the counts were normalized by dividing the counts with the corresponding effective length of the corresponding splice variant. After normalization, the relative frequency of the splice variant was calculated, and the distributions was visualized using a pie chart.

Statistical differences between different cell lines in cell growth were analyzed using analysis of variance (ANOVA), followed by Tukey’s post hoc test. Statistical significance was assumed to be meaningful if the p-value was equal to or greater than 0·05.

### 2.10 Analysis of clinical cohort and public datasets

Cox proportional hazards regression model was used to evaluate the risk of PCa death and overall death (hazard ratio, (HR)) with follow up time from initiation of ADT or ARSI treatment and death as an endpoint. Analysis was performed separately to evaluate impact of ARSI responses after ARSI initiation and overall treatment responses after initiation of ADT in patients with bicalutamide use before ARSI compared ARSI alone. Univariate analysis was adjusted with patient age at the diagnosis. Multivariate analysis was further adjusted with physical performance status, Gleason score, TNM classification, PSA level at the time of diagnosis, and use of statins, antihypertensive, or antidiabetic medication. Additional analysis was conducted including only the patients who used enzalutamide. Impact of differential treatment sequences were also illustrated using Kaplan Meier curves.

We used clinical data generated by the Cancer Dream Team consortium (SU2C data) to assess the clinical significance of the DEGs [19]. The data included samples from metastatic CRPC patients receiving standard-of-care treatment (median age at diagnosis 61·2 years). There were expression data from two different RNA-sequencing cohorts: capture cohort and polyA cohort. Cohorts were analyzed separately and the analysis was limited to the observed DEGs. Capture cohort and polyA cohort included 42 and 34 samples from ARSI-exposed patients at the time of the biopsy and 42 and 31 samples from ARSI-naïve patients, respectively. Mann-Whitney U-test was used to test the differential expression of the genes by ARSI exposure. The significance threshold was set as p-value < 0·05. R-package ggplot2 version 3·4·2 was used to generate the boxplots of expression values.

### 2.11 Data Availability

The RNA-sequencing data generated and discussed in this publication have been deposited in NCBI’s Gene Expression Omnibus (GEO) and are accessible through GEO Series accession number GSE169305 (https://www.ncbi.nlm.nih.gov/geo/query/acc.cgi?acc=GSE169305). The results published here are in part based upon data generated by the Cancer Dream Team consortium (SU2C data).

## 3. Results

### 3.1 Testosterone sensitivity

Omitting T entirely inhibited the growth of both VCaP-T and VCaP-CT. When exposed to castration level of T (0·1 nM), the growth rate of T-sensitive VCaP-T cells was significantly lower compared to the castration-resistant VCaP-CT cells (Figure 1A, p<0·0001). VCaP-CT cells grew similarly under all tested T concentrations (Figure 1A). In VCaP-BR cells, T depletion had no impact on cell growth. However, removing bicalutamide exposure increased cell growth by about 25% despite remaining under castration level of T (Figure 1B). Culturing cells with increasing concentration of T decreased the cell growth below control level. In VCaP-BRER cells, T depletion and castration levels of T slightly decreased cell growth, which was further lowered by increasing T concentration (Figure 1C). VCaP-ER cells behaved similarly to VCaP-BR cells; castration level of T first increased cell growth, however, increasing concentration of T decreased the growth below control level (Figure 1D).

**Figure 1.**
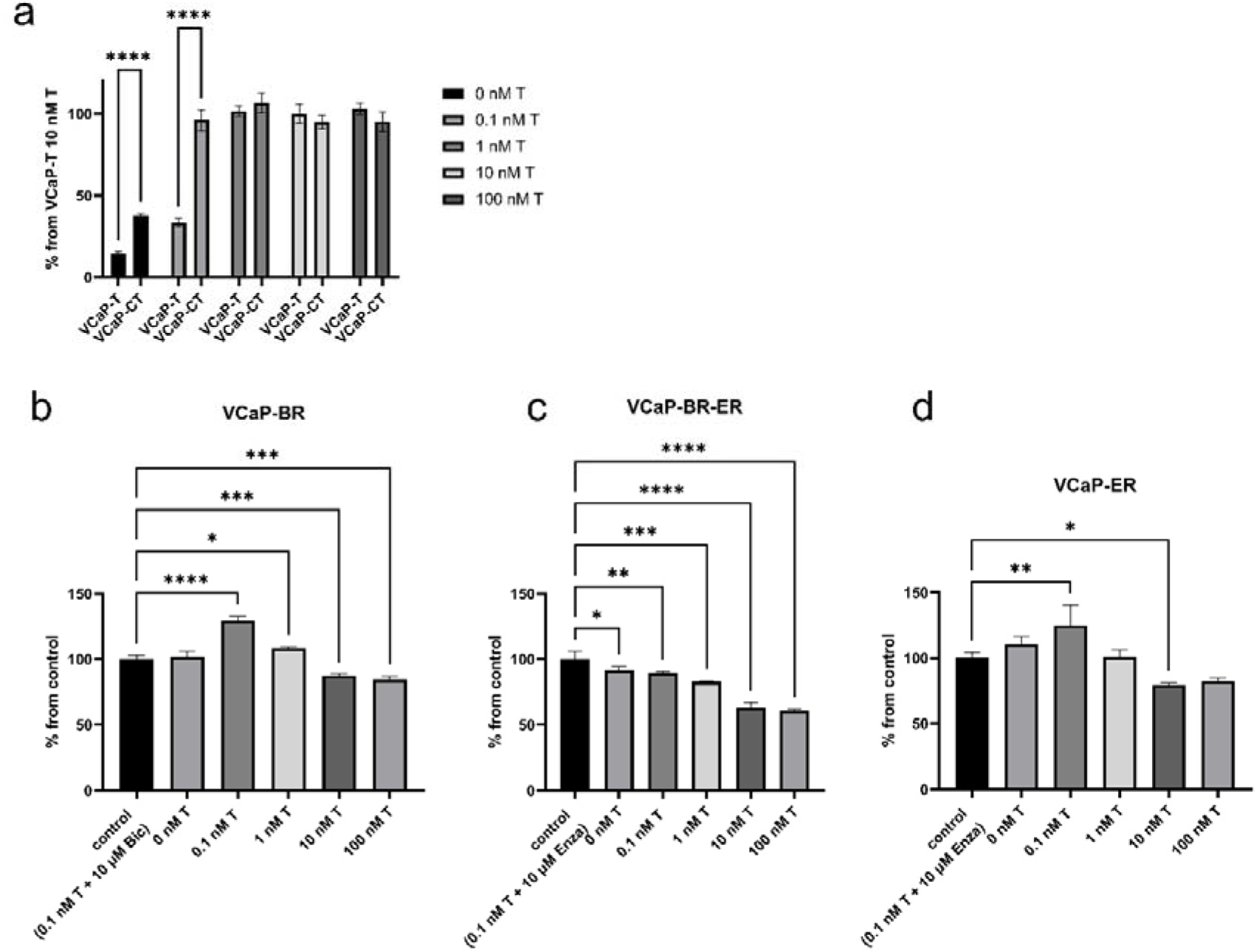
Cell growth rate diagrams under different testosterone doses. Both VCaP-T (testosterone-sensitive cells) and VCaP-CT (castration-resistant cells) showed reduced growth when cultured under T-depleted conditions compared to cells cultured under high T levels (A). The growth rate of VCaP-T cells is significantly lower than that of castration-resistant VCaP-CT cells under castrate levels of T (0·1nM) (A). The growth rates of bicalutamide-resistant VCaP-BR (B) and enzalutamide-resistant VCaP-ER cells (D) were similar at different T concentrations. T depletion had no impact on cell growth, while castration levels of T increased cell growth in both cell lines. However, increasing T concentration decreased the growth below the control level (B, D). In bicalutamide-enzalutamide-resistant VCaP-BRER cell lines, the cellular growth rate decreased with increasing testosterone concentration (C). Each treatment is presented as mean ± SD, n=3 per group. *p-value < 0·05, **p-value < 0·01, ***p-value < 0·001, ****p-value < 0·0001.

### 3.2 Transcriptomics in androgen signaling inhibitor resistant cell lines

RNA-sequencing revealed changes in transcriptomic profiles during establishment of ARSI resistance (Supplementary figure 1, Supplementary table 1). As expected, there were more DEGs in total in VCaP-ER and VCaP-BRER and they shared more DEGs compared to VCaP-T than compared to VCaP-CT (Supplementary figure 1). In both cases the shared DEGs in VCaP-ER and VCaP-BRER were regulated in the same direction.

Gene set enrichment analysis revealed similarities and differences between DEGs of different comparisons (Figure 2, Supplementary table 2-4). Androgen response was upregulated in both ARSI resistant cell lines when compared to VCaP-CT but downregulated when compared to VCaP-T (Figure 2). Interestingly, MYC targets hallmarks were positively enriched in VCaP-BRER but not in VCaP-ER. Fatty acid metabolism was upregulated in both VCaP-ER and VCaP-BRER. Hypoxia hallmark was significantly downregulated in both VCaP-ER and VCaP-BRER when compared to VCaP-T but not when compared to VCaP-CT. Unfolded protein response, which has been linked to cellular stress, was upregulated in both VCaP-ER and VCaP-BRER compared to VCaP-CT but downregulated in comparison to VCaP-T.

**Figure 2.**
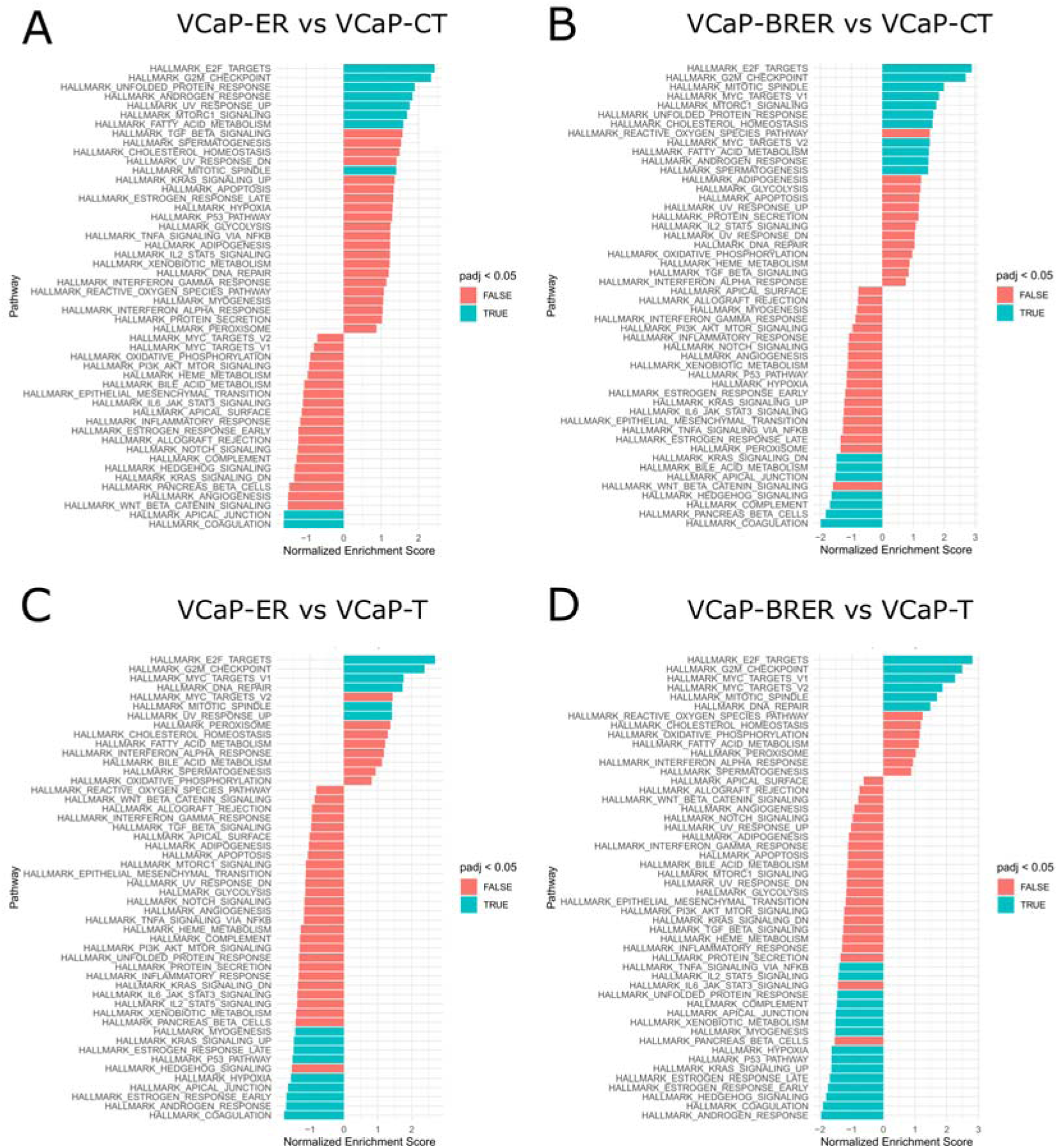
Gene set enrichment results of cancer hallmarks in different comparisons of androgen signaling inhibitor resistant cell lines to VCaP-CT and VCaP-T. Hallmarks with turquoise blocks represents significantly enriched hallmarks. Next, we focused on the DEG sections shared by or specific to VCaP-ER and VCaP-BRER either when compared to VCaP-CT or VCaP-T (Supplementary tables 5-6). A fraction of these genes has previously been reported to be AR-associated either in testosterone-dependent or testosterone-independent cell lines [20]. Over-representation analysis revealed differences and similarities in ARSI resistant cell lines (Supplementary figure 2, Supplementary table 7-10). When compared to VCaP-T the upregulated DEGs shared by VCaP-ER and VCaP-BRER were enriched in steroid biosynthesis and cellular senescence (Supplementary table 10). When compared to VCaP-CT the upregulated shared DEGs were enriched in lipid metabolism and androgen signaling while downregulated shared DEGs were enriched in exocytosis, catabolic, adhesion, and immune related terms (Supplementary figure 2A-B, Supplementary table 7). Upregulated DEGs specific to VCaP-ER were enriched in negative regulation of cell death and cancer pathways (Supplementary figure 2C). Also focal adhesion and response to unfolded proteins were enriched. Downregulated DEGs specific to VCaP-ER were enriched in cell junction and adhesion related pathways (Supplementary figure 2D). Cell division and cell cycle as well as cellular respiration related pathways were enriched among upregulated DEGs specific to VCaP-BRER (Supplementary figure 2E). Only extracellular matrix organization-related terms were enriched among downregulated DEGs specific to VCaP-BRER.

### 3.3 Transcriptomic difference between VCaP-ER and VCaP-BRER

The two enzalutamide-resistant cell lines, VCaP-BRER and VCaP-ER, display differing transcriptomic profiles (Figure 3). A side-by-side comparison exposes differences in the PCa landmark gene Kallikrein Related Peptidase 3 (*KLK3*), which shows a 3·7 log2 fold change between VCaP-BRER and VCaP-ER (Figure 3C). The androgen receptor (*AR*) was added for reference due to its importance in PCa. Gene set enrichment analysis revealed androgen signaling signature being higher in VCaP-ER than in VCaP-BRER (Figure 3D). Contrary, MYC targets hallmarks and cell cycle and checkpoint related hallmarks were higher in VCaP-BRER than in VCaP-ER. Five (*ITM2A, ID3, THBS1, PLOD2, GPX3*) of the 58 DEGs between VCaP-ER and VCaP-BRER have been previously studied to be AR-associated genes in testosterone-dependent cell line [20].

**Figure 3.**
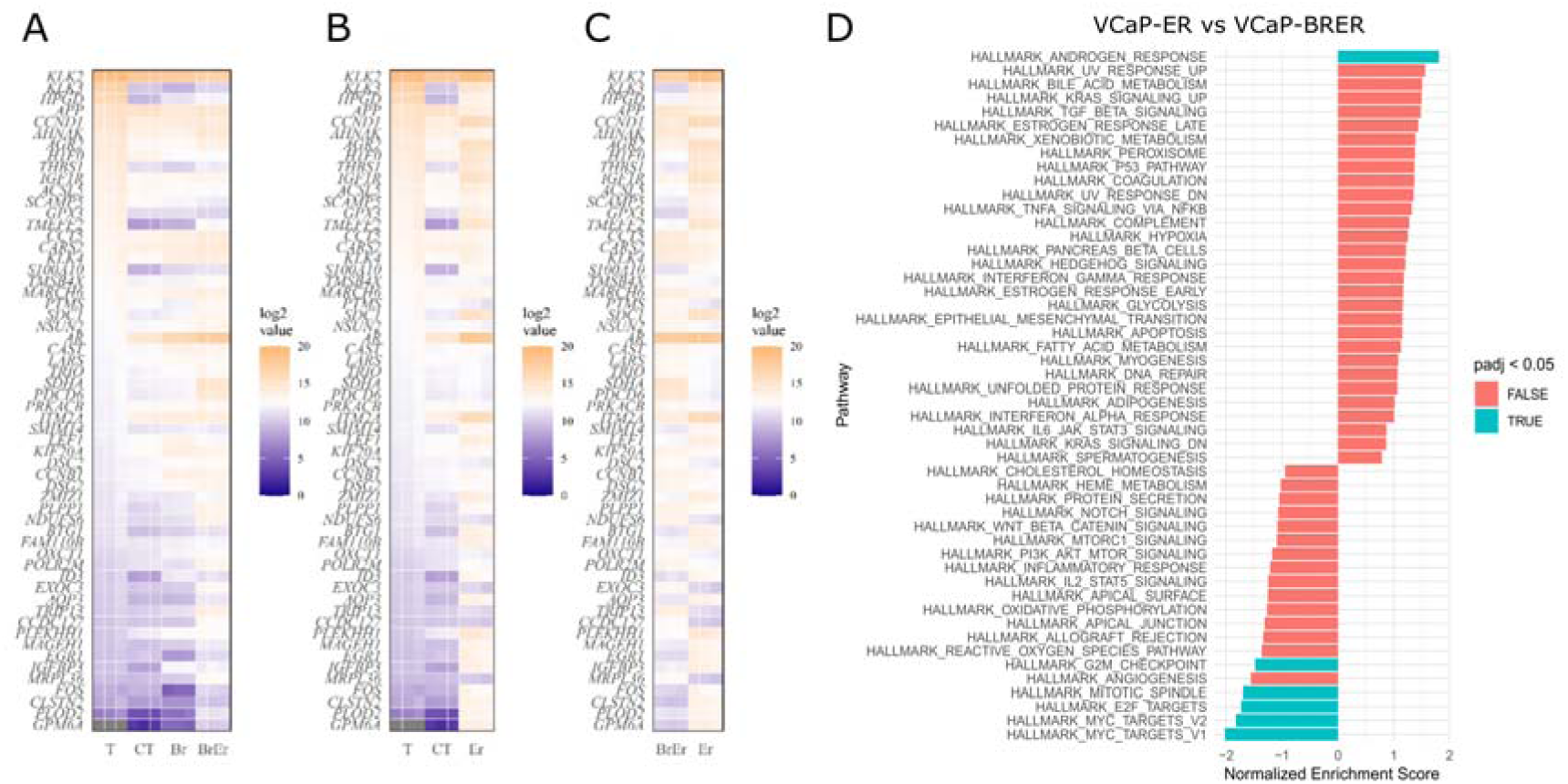
(A-C) Median ratio normalized log2 mRNA counts of differentially expressed genes (DEGs) between VCaP-BRER and VCaP-ER. The dark blue color indicates lower counts, whereas dark orange marks higher expression counts. The gene order is based on the median of the baseline sample (VCaP-T) mRNA counts. The midpoint color, white, is globally set to the median of all samples log2 transformed values. Data from three biological samples are shown. (A) Transcriptomic profile heatmap for testosterone-dependent (T), -independent (CT), bicalutamide-resistant (BR), and sequential resistance to first bicalutamide and then enzalutamide (BRER) cell lines. (B) Transcriptomic profile heatmap for testosterone-dependent (T), -independent (CT), and enzalutamide-resistant (ER) cell lines. (C) Side-by-side transcriptomic profile comparison of VCaP-BRER and VCaP-ER. (D) Gene set enrichment results of cancer hallmarks among DEGs between VCaP-ER and VCaP-BRER. Hallmarks with turquoise blocks represents significantly enriched hallmarks.

Figure 4 shows the mRNA expression levels of genes with high log2 fold-change and abundance in drug resistant cell lines. Anterior gradient 2 (*AGR2*), cyclin D1 (*CCND1*), insulin-like growth factor 1 receptor (*IGF1R*), integral membrane protein 2A (*ITM2A*), tyrosine 3-monooxygenase/tryptophan 5-monooxygenase activation protein zeta (*YWHAZ*), NK3 homeobox 1 (*NKX3-1*), and calnexin (*CANX*) are more upregulated in the VCaP-ER cell line compared to other resistant cell lines. Transmembrane protein with EGF-like and two follistatin-like domains 2 (*TMEFF2*) and N-Myc downstream-regulated 1 (*NDRG1*) are upregulated in VCaP-BRER and VCaP-ER cells compared to other cell lines, with upregulation being more evident in VCaP-ER cells. Programmed Cell Death 6 (*PDCD6*) expression is higher in the VCaP-BRER cell line compared to other cell lines. Heat shock protein 90 (*HSP90AB1*) is downregulated in VCaP-BRER and VCaP-ER cells compared to other cells. The *AR* expression is multiple-fold higher compared to VCaP-T and VCaP-CT cell lines but is indifferent between VCaP-BRER and VCaP-ER cell lines (Figure 4, Supplementary figure 3).

**Figure 4.**
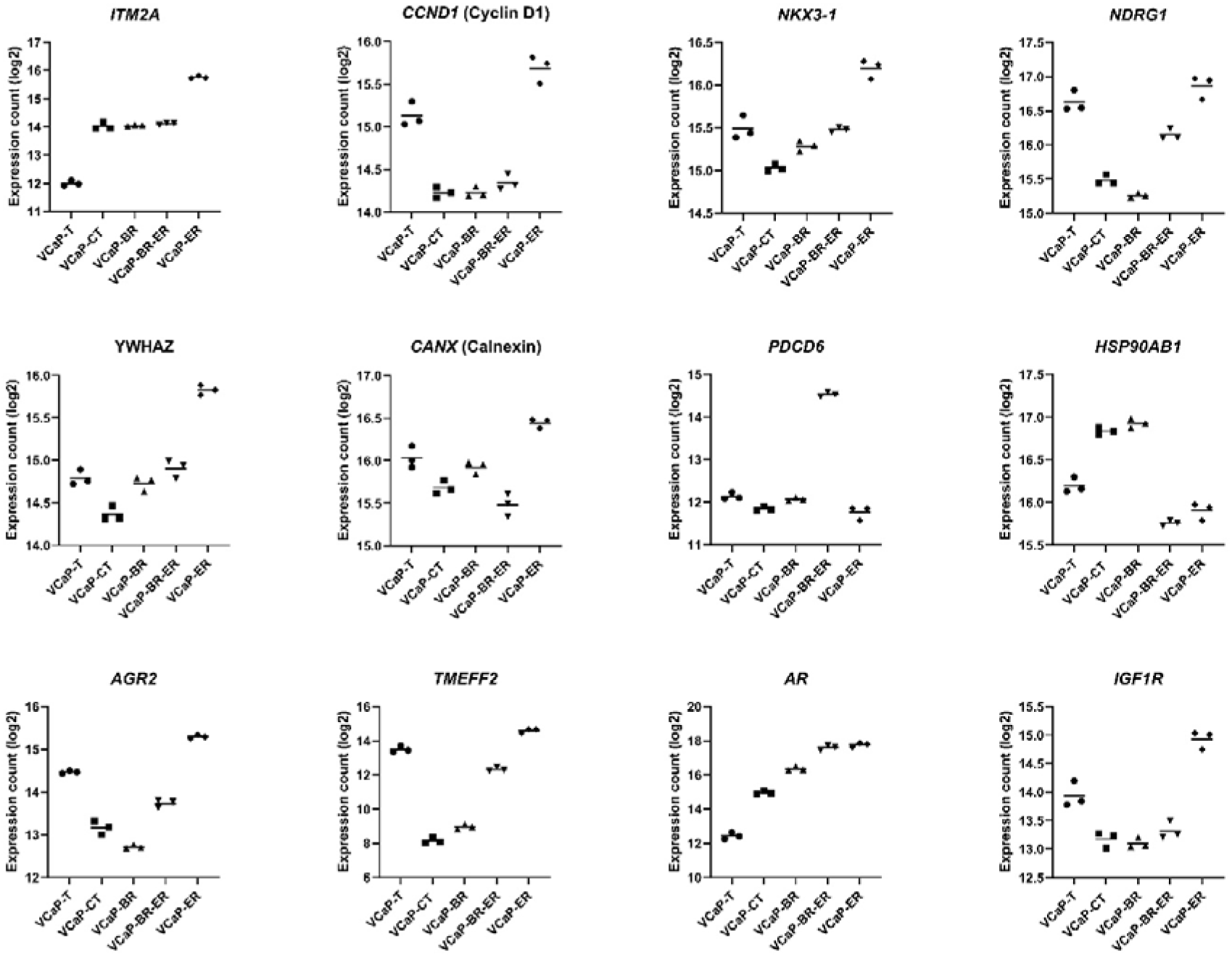
Log2 mRNA expression levels of different genes in the investigated cell lines, including integral membrane protein 2A (*ITM2A*), cyclin D1 (*CCND1*), NK3 homeobox 1 (*NKX3-1*), N-Myc downstream regulated 1 (*NDRG1*), tyrosine 3-monooxygenase/tryptophan 5-monooxygenase activation protein zeta (*YWHAZ*), calnexin (*CANX*), programmed cell death 6 (*PDCD6*), heat shock protein 90 (*HSP90AB1*), anterior gradient 2 (*AGR2*), transmembrane protein with EGF-like and two follistatin-like domains 2 (*TMEFF2*), androgen receptor (*AR*), and insulin-like growth factor receptor 1 (*IGF1R*).

### 3.4 Protein expression differences by resistance profile

NDRG1, YWHAZ, and AGR2 protein expression increased (4-5 fold, 2-3 fold, and 5-17 fold, respectively) in VCaP-BRER and VCaP-ER cell lines compared to the VCaP-CT cell line (Figure 5, Supplementary figure 4). PDCD6 expression increased in VCaP-BRER (3 fold) compared to VCaP- CT. ITM2A expression decreased (0·5 fold) in both VCaP-BRER and -ER cell lines. Nkx3·1 expression decreased only in the VCaP-ER cell line. AR and IGF1R expression increased, and TMEFF2 decreased in all cell lines compared to the VCaP-CT cell line. No difference in protein expression in cyclin D1, calnexin, and HSP90 was detected between the cell lines.

**Figure 5.**
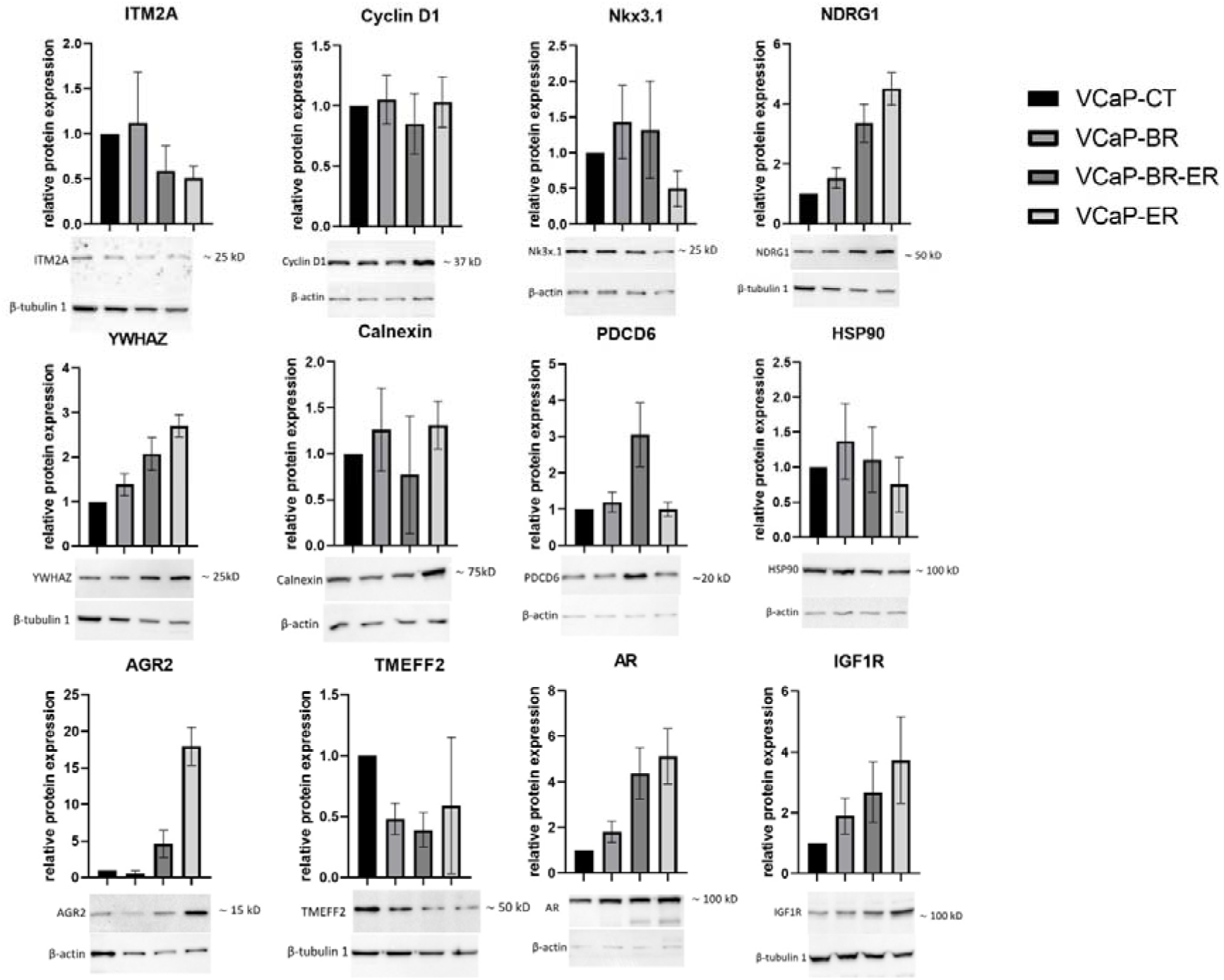
Protein concentrations (relative) of integral membrane protein 2A (ITM2A), cyclin D1, NK3 homeobox 1 (Nkx31), N-Myc downstream regulated 1 (NDRG1), tyrosine 3-monooxygenase/tryptophan 5-monooxygenase activation protein zeta (YWHAZ), calnexin, programmed cell death protein 6 (PDCD6), heat shock protein 90 (HSP90), anterior gradient 2 (AGR2), transmembrane protein with EGF-like and two follistatin-like domains 2 (TMEFF2), androgen receptor (AR), and insulin-like growth factor 1 receptor (IGF1R) for VCaP-CT, VCaP-BR, VCaP-BRER, and VCaP-ER cell lines. The medium gray bar represents the bicalutamide-resistant VCaP-BR cell line, the dark gray bar represents the bicalutamide- and enzalutamide-resistant VCaP-BRER cell line, and the light gray bar represents the enzalutamide-resistant VCaP-ER cell line. The protein concentrations are relative to the castration-resistant VCaP-CT cell line (black bar). Examples of cropped western blots are presented and full blots of all replicates are included in Supplementary figure 4.

A comparison between the transcriptome and protein expression shows that changes in AGR2, IGF1R, PDCD6, YWHAZ, and NDRG1 are parallel in both (Figure 4 and 5). On the contrary, ITM2A and TMEFF2 mRNA upregulation reflects as decreased protein concentration. CCND1 upregulation is not followed by an increased CCND1 protein concentration.

### 3.5 Splice variant differences by resistance profile

*AGR2* and *PDCD6* exhibit alternative splicing in VCaP-BRER cells, as compared to VCaP-T, VCaP-CT, and VCaP-BR cells (Supplementary figure 5). However, in VCaP-ER cells, where enzalutamide was directly given without the preceding bicalutamide, the *AGR2* and *PDCD6* splice variant profile did not change in comparison to VCaP-T and VCaP-CT. Otherwise, the cells did not demonstrate significant changes in their splice variant profiles. VCaP-BRER and VCaP-ER exhibit increased relative expression of *AR* splice variant AR-004, which corresponds to AR-V7, whereas other cell types showed no significant differences.

### 3.6 Phenotype of androgen signaling inhibitor resistant cell lines

All cells were sensitive to docetaxel at concentrations ranging from 1 nM to 10 nM. However, sensitivity to docetaxel differed between the cell lines (Figure 6). The VCaP-BRER cells showed the highest resistance to docetaxel exposure (around 50% of cells viable after 10 nM dose) and VCaP-BR the lowest (around 25% of cells viable after 10 nM dose). VCaP-CT and VCaP-ER showed similar responses to docetaxel (around 40% of cells viable after 10 nM dose).

**Figure 6.**
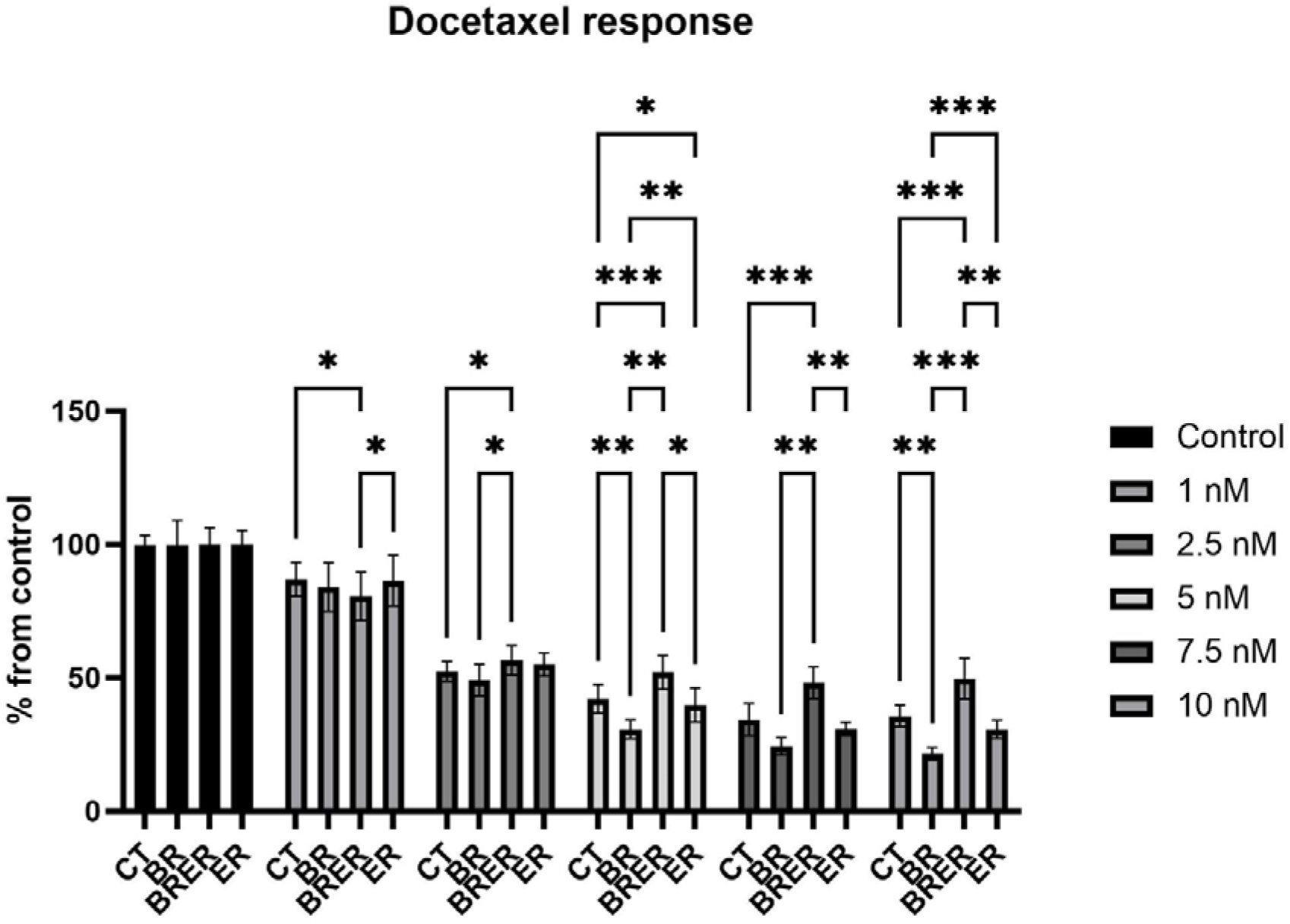
Relative cell viability under docetaxel exposure at increasing doses 1 nM to 10 nM. All cells were sensitive to docetaxel exposure, however, the cellular response slightly differed between the cell lines. The VCaP cells resistant to both enzalutamide and bicalutamide (ERBR) cells display highest resistance and VCaP cells resistant only to bicalutamide (BR) cells show lowest resistance to docetaxel exposure. Castration resistant cells (CT) and enzalutamide resistant cells (ER) showed similar responses to docetaxel in all investigated concentrations.

Brightfield images of each cell line are shown in Figure 7. Androgen-dependent (Figure 7A) and -independent (Figure 7B) VCaP cells grew in tightly arranged cell batches. The former was characterized by a smooth outline, while the latter showed occasional short membrane protrusions, extent and number of which increased with the development of antiandrogen resistance (Figure 7C-E). While bicalutamide-resistant cells grew in close contact with each other (Figure 7C), the development of enzalutamide resistance (Figure 7D-E) was accompanied by a more diffuse cellular arrangement.

**Figure 7.**
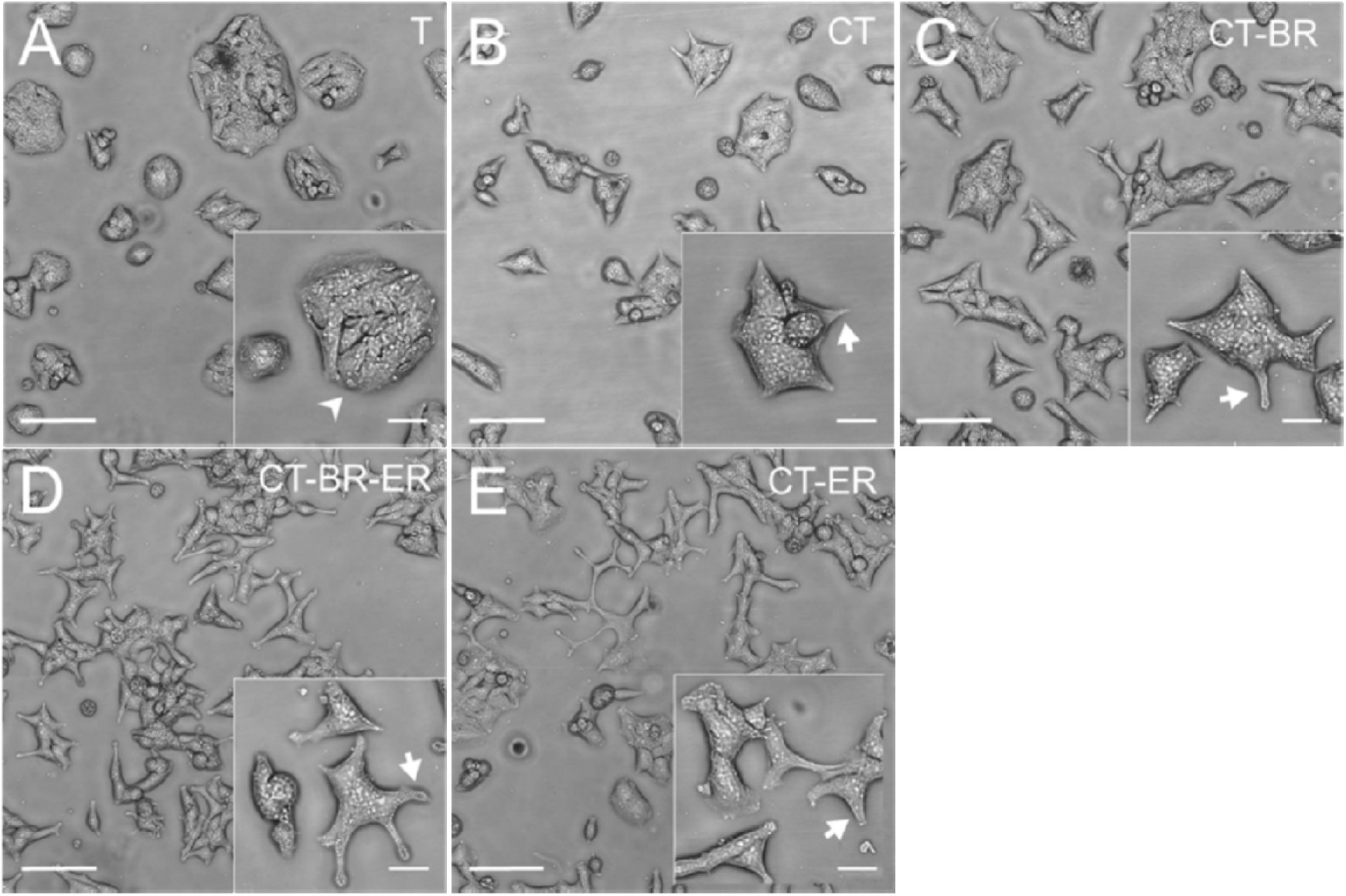
Growth characteristics and morphology of VCaP cell lines with different sensitivity to testosterone (A-B) and drug resistances (C-E). The cell lines were seeded at equal densities in T25 culture bottles and maintained for 6d in culture prior to microscopic analysis on a Nikon Ti-S microscope. (A) Androgen-dependent VCaP-T cells grow in tight cell batches with a smooth outline (arrowhead in inset). (B) Androgen-independent VCaP-CT cells exhibit tightly arranged cell groups with occasional short membrane protrusions (arrow in inset). An increase in the number and extent of protrusions is seen in cells displaying antiandrogen-resistance (C-E; arrow in insets). Bicalutamide-resistant VCaP-BR cells (C) grow in close contact with each other whereas in both enzalutamide-resistant cell lines VCaP-BRER (D) and VCaP-ER (E) a more diffuse growth pattern predominates. Bars 100 µm; in insets 25 µm.

### 3.7 Evaluation of clinical relevance

Population characteristics are presented in Supplementary table 11. The cohort consisted of 493 males with castration-resistant prostate cancer. The majority of patients had enzalutamide as their form of second generation ARSI therapy (85,6%). The group of patients who had used bicalutamide before second generation ARSI therapy consisted of 201 males with a median (IQR) age of 78 (72-84) years while bicalutamide non-users consisted of 292 males with a median (IQR) age of 75 (70-80) years (p<0.05 between the groups). In the group of bicalutamide users, 16,4 % of patients had a locally spread tumor (N1), and 27,9 % of patients had a metastatic tumor (M1) while bicalutamide non-users, 33,2 % had a locally spread tumor and 52,1 % had a metastatic tumor at the time of diagnosis (p<0.05 for both).

Based on our age-adjusted analysis of ARSI treatment response, the risk of PCa-specific death was elevated in the group of antecedent bicalutamide use compared to ARSI use alone (HR, 1.75; 95 % CI, 1.38-2.22). Similarly, the risk of overall death was elevated in bicalutamide user group (HR, 1.76; 95 % CI, 1.41-2.21). Figure 8A and 8B illustrates the trend of PCa-specific death and overall death, respectively, with Kaplan-Meier curves.

**Figure 8.**
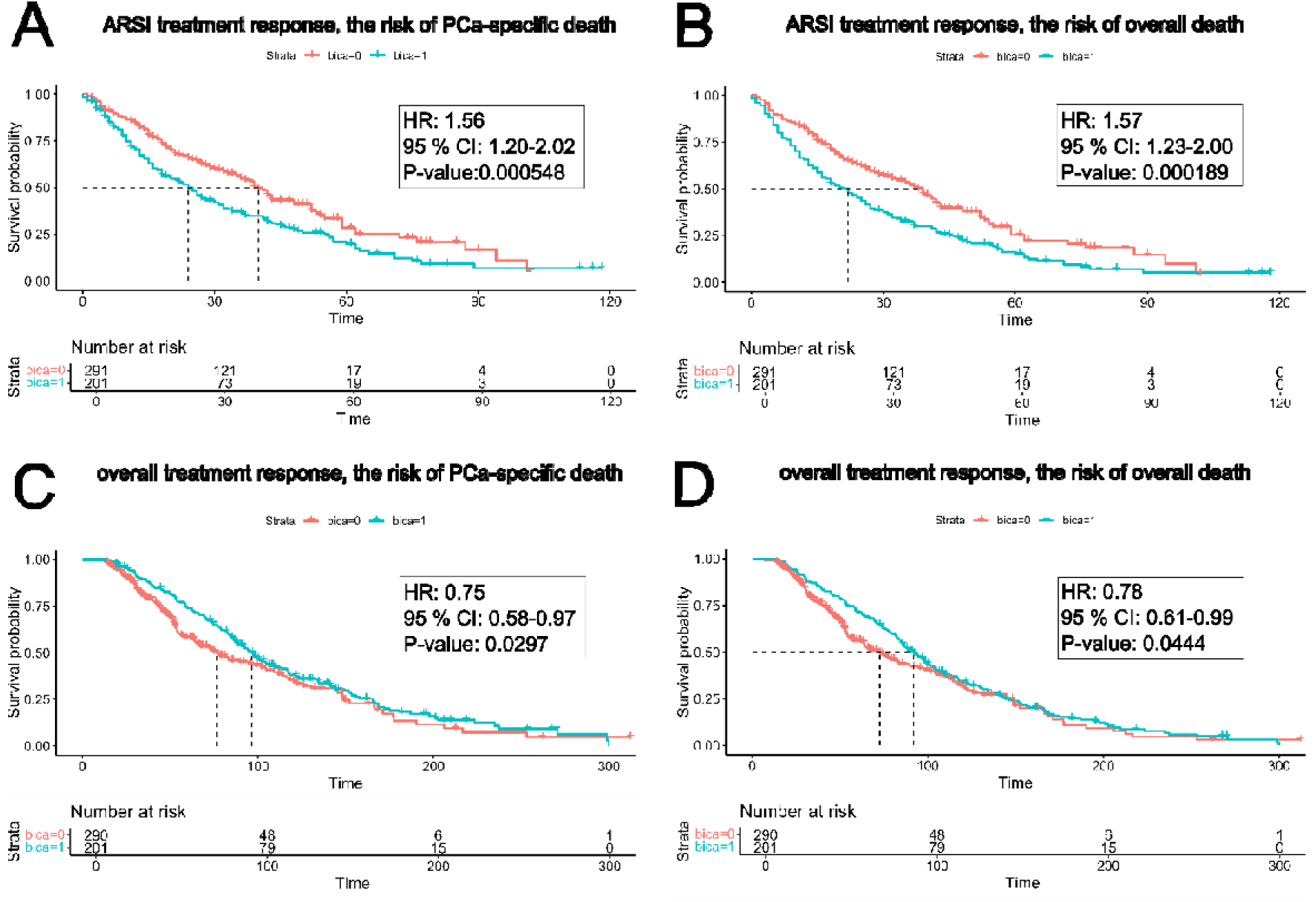
Kaplan-Meier curves presenting the risk of PCa-specific death in ARSI treatment (A), the risk of overall death in ARSI treatment (B), the risk of PCa-specific death overall in all treatments of adanced PCa (C) and the risk of overall death overall in all treatments of adanced PCa (D). Patients are stratified based on whether they have received bicalutamide before second generation ARSI therapy (bica=1) or not (bica=0). Hazard ratio (HR), confidence interval (CI) and p-value from multivariable-adjusted Cox regression analysis are presented along with the Kaplan-Meier curves.

In our multivariable-adjusted analysis evaluating ARSI treatment responses, bicalutamide use before second generation ARSI therapy was associated with increased risk of both PCa-spesific death (HR, 1.56; 95 % CI, 1.20-2.02) and overall death (HR, 1.57; 95 % CI, 1.23-2.00). In the multivariable-adjusted analysis of only enzalutamide users, antecedent use of bicalutamide was associated with increased risk of both PCa-spesific death (HR, 1.45; 95 % CI, 1.11-1.90) and overall death (HR, 1.44; 95 % CI, 1.11-1.85).

When analysis was done to evaluate overall response to all treatments for advanced PCa, bicalutamide use before ARSI therapy was not associated with overall death (HR, 0.91; 95 % CI, 0.72-1.13) or PCa-spesific death (HR, 0.88; 95 % CI, 0.69-1.11) using age-adjusted analysis. In multivariable-adjusted analysis evaluating overall response to all treatments, bicalutamide use before ARSI therapy was associated with mildly decreased risk of both PCa-specific death (HR, 0.75; 95 % CI, 0.58-0.97) and overall death (HR, 0.78; 95 % CI, 0.61-0.99). When evaluating the overall response to all treatments with multivariable-adjusted analysis among only enzalutamide users the risk of PCa-specific death (HR, 0.76; 95 % CI, 0.58-0.98) and overall death (HR, 0.78; 95 % CI, 0.61-1.00) were similar to the risk among all patients. Figure 8C and 8D illustrates the trend of PCa-specific death and overall death, respectively, with Kaplan-Meier curves.

Further we analyzed the expression of the DEGs in our cell lines also in public clinical data including metastatic castration-resistant prostate cancer samples (SU2C data) [19]. We compared the expression of DEGs between ARSI-exposed and ARSI-naïve PCa patients in two RNA-sequencing cohorts (p < 0·05). *AR* was upregulated among the ARSI-exposed in both sequencing cohorts (Figure 9A-B). Also, among the DEGs between VCaP-ER and VCaP-BRER the genes *BTG1* and *SDC1* were upregulated by ARSI exposure in capture cohort (Figure 9C-D). In addition, among DEGs between VCaP-ER and VCaP-CT, *FKBP5* was upregulated in both cohorts (Figure 9E-F), while *PTPN13* and *TBC1D16* were downregulated among ARSI-exposed patients in capture cohort (Figure 9G-H). *ZMIZ1* was upregulated by ARSI-treatment in polyA cohort (Figure 9I). Among DEGs shared by VCaP-ER and VCaP-BRER compared to VCaP-CT, *APOD* and *SDC1* were upregulated (Figure 9J and D) and *ENDOD1* was downregulated among ARSI-exposed patients in capture cohort (Figure 9K). The directions of the expression differences were in coherence between the cell lines and clinical samples.

**Figure 9.**
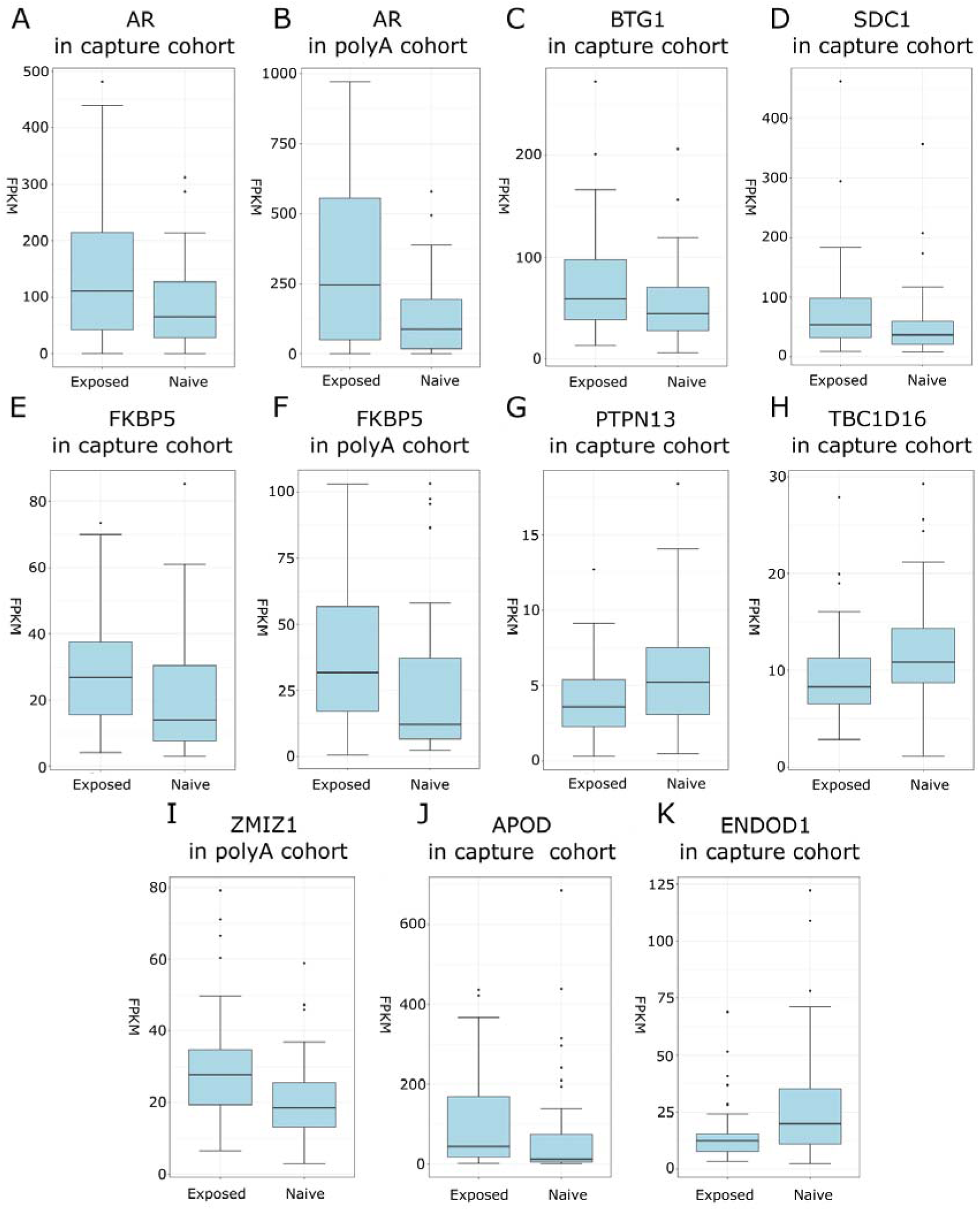
Boxplots of the expressions (FPKM) in metastatic castration-resistant prostate cancer patients (PRAD SU2C 2019 data) of the genes (A-K) that showed differential expression between patients that have been exposed to androgen signaling inhibitors and patients that are naïve to androgen signaling inhibitors in either or both of the sequencing cohorts (capture cohort or/and polyA cohort).

## 4. Discussion

Previous studies have demonstrated that enzalutamide is more effective in treating castration-resistant PCa than bicalutamide or other forms of first generation antiandrogens [21]. This would suggest that initiating second generation ARSI therapy without antecedent bicalutamide therapy leads to a longer progression-free period and overall survival time in CRPC patients. Indeed, the difference in PCa survival caused by treatment sequencing was observed in our clinical cohort; use of bicalutamide before second generation ARSI therapy was associated with increased risk of both PCa-specific and overall death when compared to ARSI alone. However, when we analyzed effectiveness of all advanced PCa treatments between the groups we did not observe differences between PCa-specific or overall mortality or the difference was slightly opposite.

Certain similarities in the mechanism of action can be found between bicalutamide and ARSI, and therefore, the decreased effectiveness of post-bicalutamide ARSI could be due to pre-existing resistance towards ARSI caused by preceding use of bicalutamide. This notion is supported by the fact that overall responses to the treatment of advanced PCa was similar.

Acquiring androgen-independence and resistance to enzalutamide and/or bicalutamide induces significant phenotypic and transcriptomic changes in PCa cells. We show that these changes depend on both the drug used and the sequence of prior antiandrogen exposures. Sequential resistance to bicalutamide and enzalutamide leads to distinct transcriptomic and phenotypic profile compared to direct enzalutamide resistance, which correlate with different docetaxel sensitivity. Thus, treatment sequencing may influence response to subsequent therapies. Several DEGs were linked to ARSI exposure, providing potential targets to improve advanced PCa therapy.

The castration-resistant VCaP-CT cell line required only castrate levels of testosterone (T) for normal growth, while drug-resistant lines VCaP-BR, VCaP-BRER, and VCaP-ER did not require T. In these resistant cells, high T concentrations even inhibited growth. Elevated androgen receptor (AR) expression, both at mRNA and protein levels, was observed in these lines, similar to clinical PCa resistance [22,23]. Gene enrichment analysis showed upregulation of androgen response pathways in resistant cells compared to VCaP-CT. These findings suggest that despite AR signaling inhibition, resistant cells activate AR signaling, and excessive T may lead to AR overstimulation, explaining growth inhibition.

Bipolar androgen therapy (BAT) has been tested in mCRPC patients resistant to ARSIs [24–26]. In BAT, patients receive monthly testosterone injections while on ADT, cycling serum testosterone between high and castrate levels. This approach yields similar clinical responses to enzalutamide and may re-sensitize enzalutamide-resistant patients. Future studies will assess whether BAT re-sensitizes our cell models to ARSIs and its impact on cell transcriptomes and phenotypes.

Treatment of castration-resistant VCaP cells with enzalutamide alone resulted in a distinct transcriptomic profile compared to sequential bicalutamide and enzalutamide treatment. Both treatments upregulated androgen signaling, with more pronounced response with direct enzalutamide resistance. Additionally, the unfolded protein response was upregulated in both cases, with specific enrichment in direct enzalutamide resistance. Thus, unfolded protein response is partly regulated by distinct genes in direct and sequential enzalutamide resistance. MYC target pathways were upregulated only in sequential resistance, while direct resistance upregulated negative regulation of cell death. Sequential resistance, in contrast, upregulated pathways related to cell cycle, division, and respiration. These differences suggest varying mechanisms of cancer cell proliferation and progression. Additionally, morphological changes were observed, reflecting the acquired drug resistance.

Our findings show that different sequences of antiandrogenic treatments create distinct selection pressures on PCa cells, leading to varied mechanisms of enzalutamide resistance. These differences in transcriptomic profiles and morphology also affected responses to docetaxel. Bicalutamide-resistant cells showed the lowest cell viability to docetaxel, while sequentially resistant cells exhibited higher resistance. This suggests that treatment sequencing may enhance susceptibility to subsequent therapies like docetaxel, with potential clinical implications.

Higher expression of AGR2, CCND1, IGF1R, ITM2A, CCND1, NDRG1, and TMEFF2 was observed after direct resistance compared to sequential enzalutamide resistance. PDCD6 displayed the opposite trend, being only upregulated when the cells were resistant to bicalutamide before the enzalutamide resistance. The parallel transcriptomic and proteomic changes in AGR2, IGF1R, and PDCD6 indicate their activity in the cell. Moreover, AGR2 and PDCD6 displayed alternative splicing profiles in the bicalutamide-enzalutamide-resistant cells, compared to other cell types, further indicating the importance of these genes in the cellular adaptation mechanism against AR inhibitors. Based on literature, AGR2 has a role in cell migration and metastasis and acts as a p53 inhibitor, IGF1R has role in transformation events and acts as an anti-apoptotic agent and PDCD6 is calcium-binding protein and participates in programmed cell death [27]. *IGF1R* has been associated with PCa progression, while *YWHAZ, NDRG1,* and *AGR2* with aggressive PCa [28–31]. *IGF1R* has also been associated with treatment resistance in PCa and other solid cancers [32–34]. Many DEGs observed in our study have previously been related to regulation of cell proliferation, apoptosis, and adhesion during cancer progression [35–37].

To further evaluate the clinical significance of our results we analyzed the association to ARSI exposure of the observed DEGs in PRAD SU2C data. Multiple of the DEGs observed in our cell lines were differentially expressed between patients exposed to ARSIs and those naïve to androgen signaling inhibitor and a part of these were related to lipid metabolism (*APOD, PTPN13, SDC1*). For example, Apolipoprotein *APOD* is dysregulated in PCa and has previously been found to be a marker for PCa [38]. We found *APOD* to be upregulated during formation of both resistances as well as in ARSI-exposed patients in comparison to ARSI-naïve patients. Also gene enrichment analysis revealed lipid metabolism being upregulated during both direct and sequential enzalutamide resistance, emphasizing its importance for PCa progression and also for antiandrogen resistance.

*ENDOD1,* which was downregulated during formation of both resistances, has been previously found to be downregulated in high-grade and metastatic PCa compared to low-grade PCa in cell line models [39,40]. We further showed that its expression was downregulated also in ARSI-exposed patients in comparison to ARSI-naïve patients. AR regulated *FKBP5,* which has been previously linked to chemoresistance [41], was upregulated in ARSI-exposed patients in comparison to ARSI-naïve patients and during the development of direct enzalutamide resistance in cell line model. Also *SDC1* has been previously linked to chemoresistance, found to be more expressed in aggressive PCa and associated to unfavorable prognosis [42,43]. Here *SDC1* was upregulated during formation of both resistances, although more upregulated in direct resistance to enzalutamide compared to sequential resistance, and among ARSI-exposed patients compared to ARSI-naïve patients. These findings shows that our model is in line with previous studies and extends the previous results regarding these genes.

*PTPN13* has been studied to be downregulated and having tumor suppressing role in multiple cancers but has not been studied in PCa [44–47]. Here it was specifically downregulated during formation of direct enzalutamide resistance and among ARSI-exposed patients compared to ARSI-naïve patients. Also *BTG1* has a tumor suppressing role in several cancer types but has not been studied in PCa [48]. We found *BTG1* to be upregulated in direct enzalutamide resistance compared to sequential resistance as well as in ARSI-exposed patients compared to ARSI-naïve patients. These findings associate these cancer genes also to prostate cancer.

One of the main strengths of this study is that all cell-lines used were grown without acceleration to mimic naturally occurring transcriptomic changes. The RNA-seq data was complemented with protein concentration measurements to validate the biological relevance of the differentially expressed genes, which benefits the study. Moreover, to bring our results into a biological context we evaluated phenotype characteristics of the cell lines in terms of docetaxel resistance and cell morphology. We also used clinical data to emphasize the clinical significance. The main weakness of this study is the lack of *in vivo* validation. Moreover, only one type of PCa cells (VCaP) was used, thus validation will be needed from other PCa cell lines. The clinical cohort study is limited by its retrospective and non-randomized nature and, therefore, the possibility of residual bias remained despite adjusting for confounding factors.

In conclusion, the transcriptome and proteome of castration-resistant PCa cell lines depend on which drugs they have developed resistance, and on the sequence in which the resistance has developed. Moreover, they modulate docetaxel response and cellular morphology. This suggests, that by selecting the treatment lines, it may be possible to drive the PCa transcriptome to a certain direction. Even though, sequential treatment of bicalutamide and ARSI did not impact on overall benefit of all treatments for advanced PCa, it impacted on benefit of ARSI treatment after formation of CRPC. A novel idea is to find, select, and sequence the treatments to affect PCa cell transcriptome to become more sensitive to the next lines of treatment.

## Acknowledgements

The authors thank Niina Ikonen and Kati Rouhento for excellent technical assistance. This study was supported by Finnish Functional Genomics Centre, University of Turku and Åbo Akademi and Biocenter Finland. The authors wish to acknowledge CSC – IT Center for Science, Finland, for computational resources. The results published here are in part based upon data generated by the Cancer Dream Team consortium. The study has been funded by Cancer Foundation Finland, Sigrid Jusélius Foundation, Academy of Finland, The Finnish Cultural Foundation and Emil Aaltonen Foundation. ChatGPT was partly used to assist the summarization of the text.

## Author Contributions

Reetta Nätkin: Conceptualization, Data curation, Formal Analysis, Funding acquisition, Investigation, Methodology, Software, Validation, Visualization, Writing – original draft, Writing – review & editing. Milja I. Paloneva: Conceptualization, Data curation, Formal Analysis, Methodology, Validation, Writing – original draft, Writing – review & editing. Paavo V. H. Raittinen: Conceptualization, Data curation, Formal Analysis, Investigation, Methodology, Software, Validation, Visualization, Writing – original draft, Writing – review & editing. Heimo Syvälä: Conceptualization, Investigation, Methodology, Validation, Visualization, Writing – review & editing. Merja Bläuer: Conceptualization, Investigation, Methodology, Visualization, Writing – review & editing. Teuvo L. J. Tammela: Conceptualization, Funding acquisition, Investigation, Project administration, Resources, Supervision, Writing – review & editing. Matti Nykter: Conceptualization, Investigation, Resources, Supervision, Writing – review & editing. Pauliina Ilmonen: Conceptualization, Investigation, Resources, Supervision, Writing – review & editing. Aino Siltari: Conceptualization, Data curation, Formal Analysis, Funding acquisition, Investigation, Methodology, Project administration, Supervision, Validation, Visualization, Writing – original draft, Writing – review & editing. Teemu J. Murtola: Conceptualization, Funding acquisition, Investigation, Project administration, Resources, Supervision, Writing – original draft, Writing – review & editing

## Ethics Statement

Patient data were collected from the Wellbeing Services County of Pirkanmaa database with research permit R24223.

