## Supplementary Figures for "Phenotypic and transcriptomic characterization of bicalutamide and enzalutamide resistance in castration-resistant prostate cancer cells"

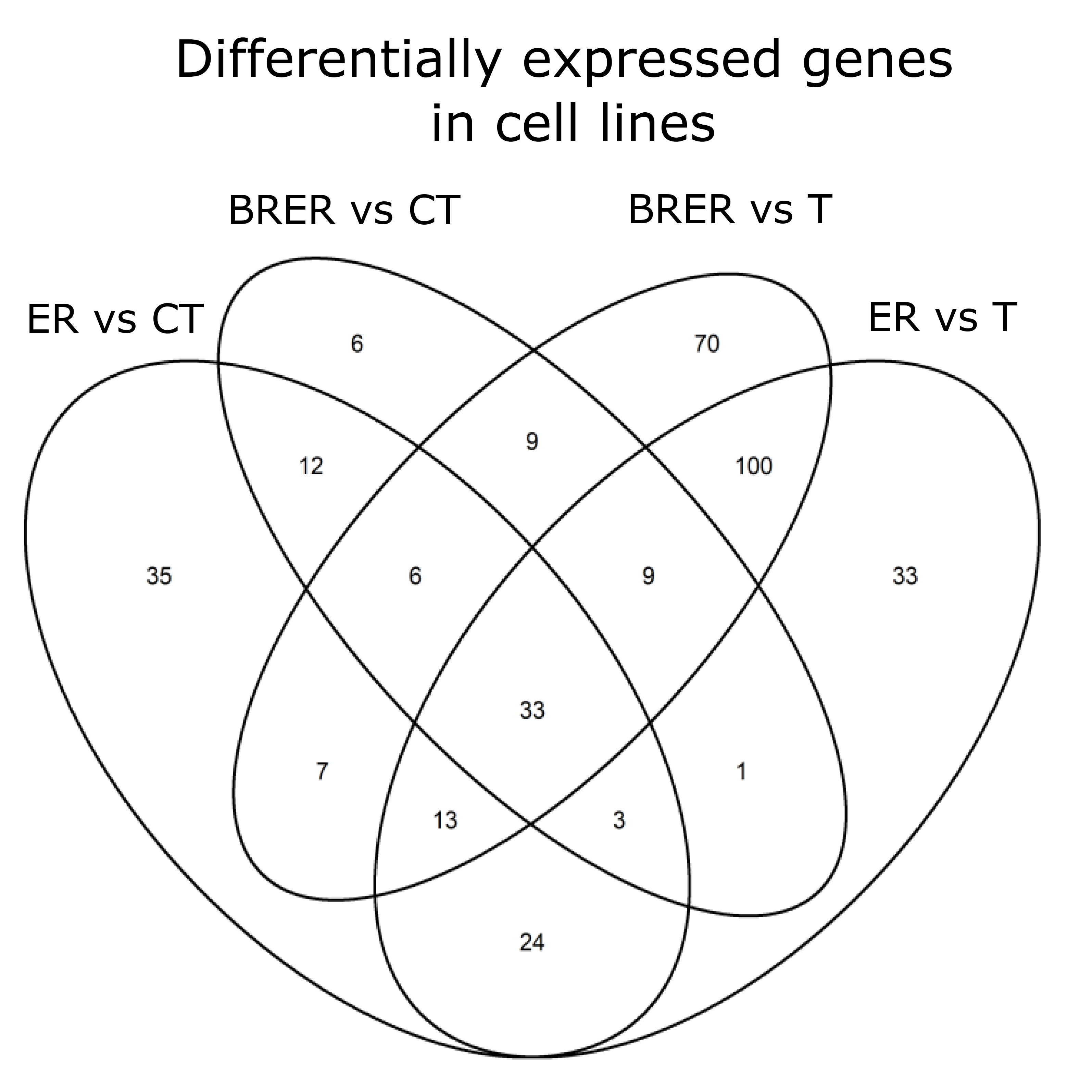

**Supplementary figure 1.** Venn diagram of shared and comparison specific differentially expressed genes (DEGs). All shared DEGs were regulated in same direction.

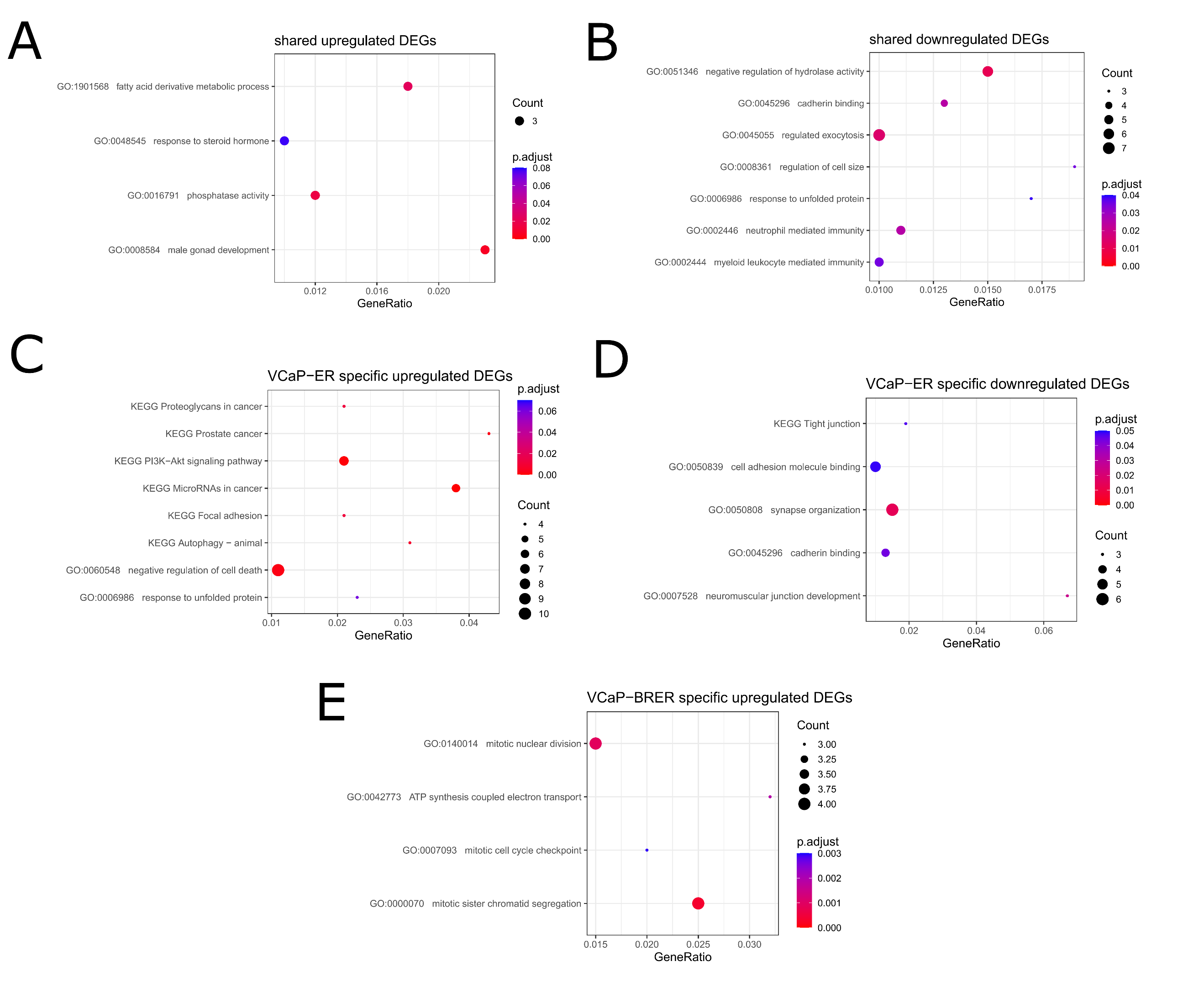

**Supplementary figure 2.** Selected results (see Methods) of over-representation analysis among DEGs shared by or specific to VCaP-ER or VCaP-BRER when compared to VCaP-CT. Downregulated DEGs specific to VCaP-BRER are not presented due to small total amount of enriched terms. Count describes the amount of significant genes found from the term/pathway and GeneRatio describes the fraction of significant genes from the total amount of genes annotated to the term/pathway.

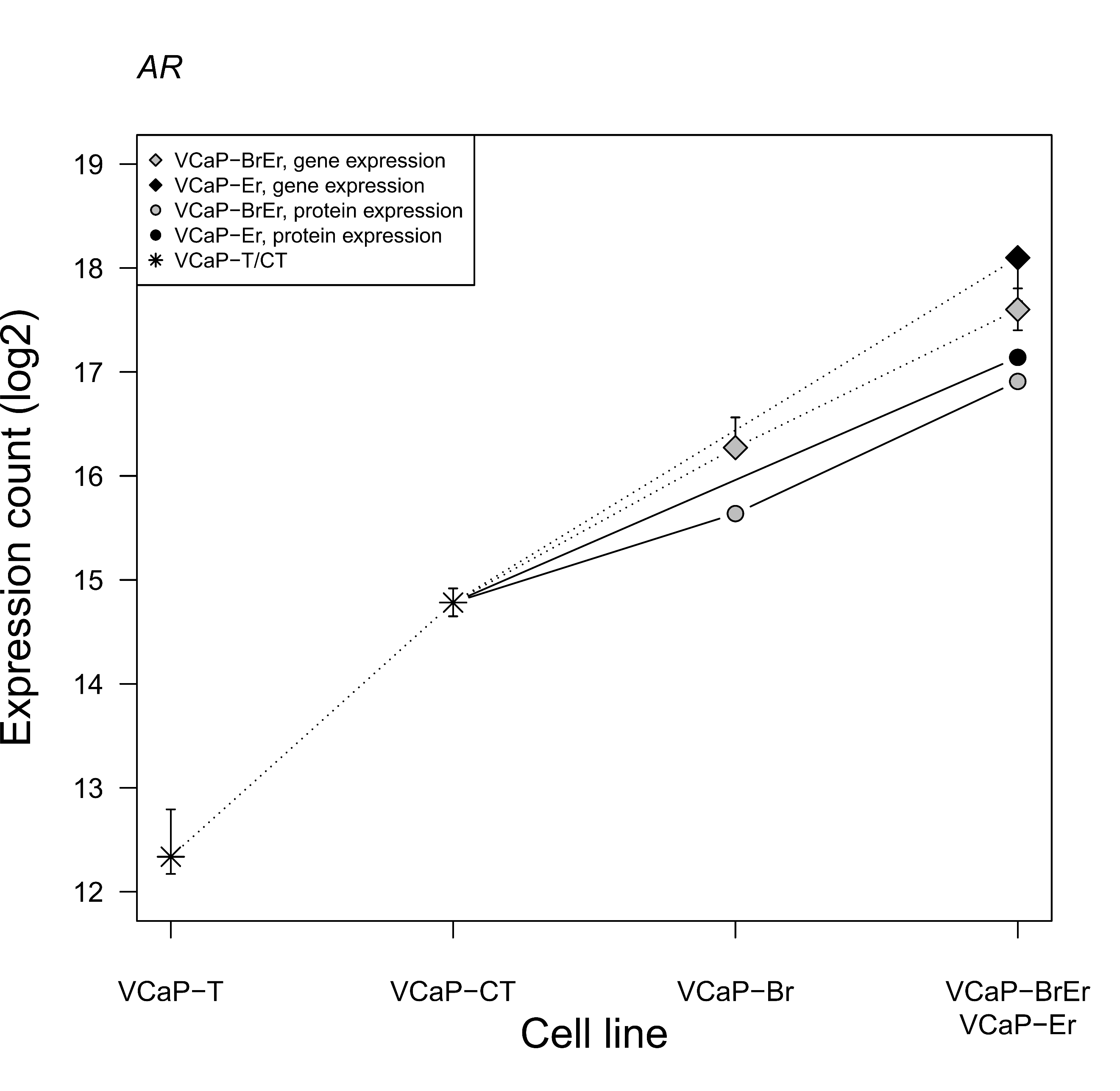

Supplementary figure 3. Cell and protein expression curves for AR. The circles filled with grey and black, connected with a solid line, represent the protein expression. The grey and black diamonds, connected with a dashed line, represent the gene expression. To preserve image clarity, the stars represent the gene expression for VCaP-T and VCaP-CT. The VCaP-CT protein expression is scaled based on the VCaP-CT gene expression to enable comparison between protein and gene expression curves. Parallel black and dashed lines remark parallel protein and gene expression whereas divergent lines indicate dissimilar protein and gene expression profiles. AR gene and protein expression does not show significant difference between VCaP-BRER and VCaP-BR.

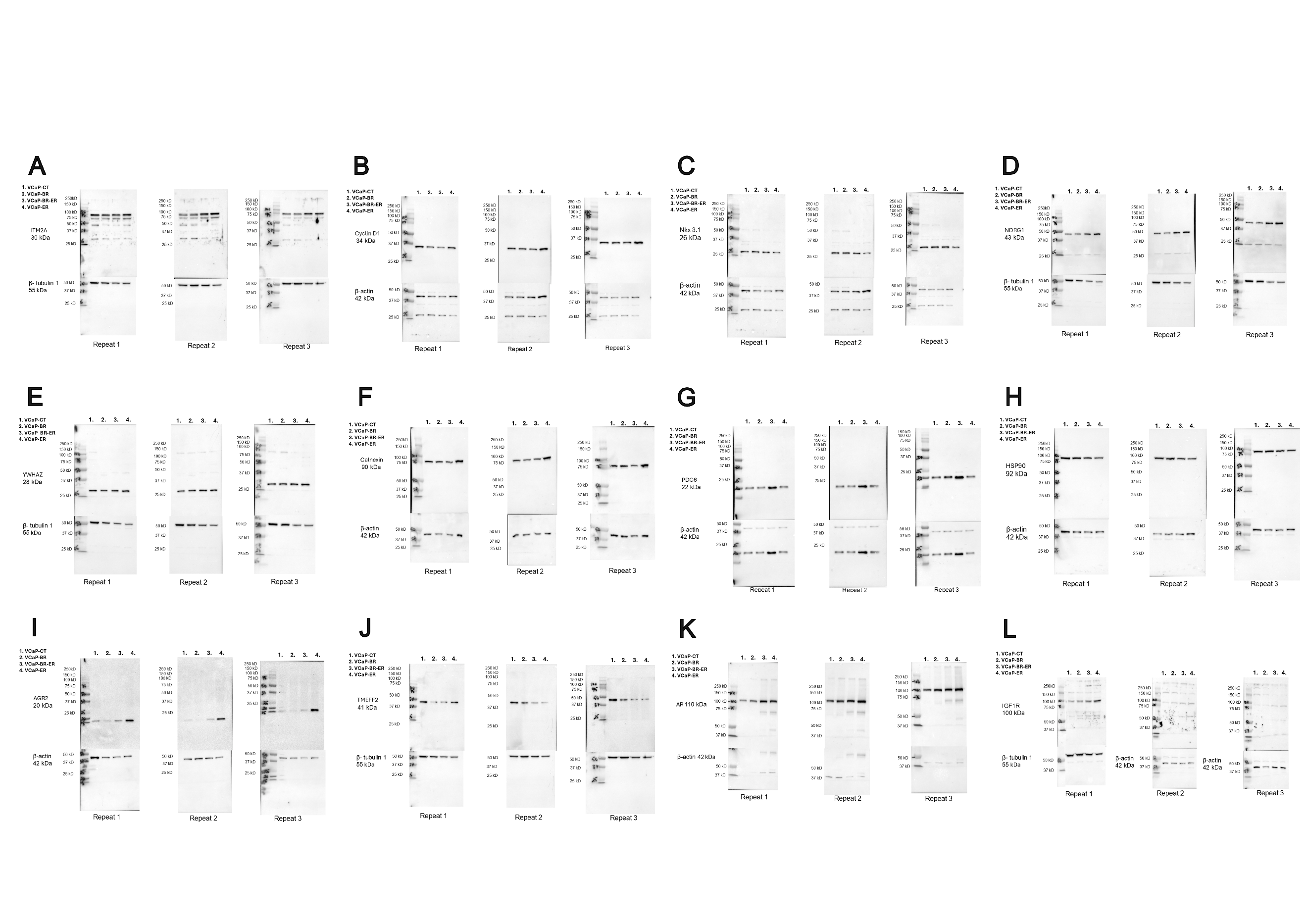

**Suplementary Figure 4.** Figure represents original whole western blot analyses of each protein shown in Fig 5. Barplots in Fig 5 presents statistics from all three (3) replicates and one representative western blot of each protein has been selected to be shown in Fig 5. Panel A shows western blot analyses for protein ITM2A, Panel B for protein Cyclin D1, Panel C for protein Nkx3.1, Panel D for protein NDRG1, Panel E for protein YWHAZ, Panel F for protein Calnexin, Panel G for protein PDCD6, Panel H for protein HSP90, Panel I for protein AGR2, Panel J for protein TMEFF2, Panel K for protein AR, Panel L for protein IGF1R.

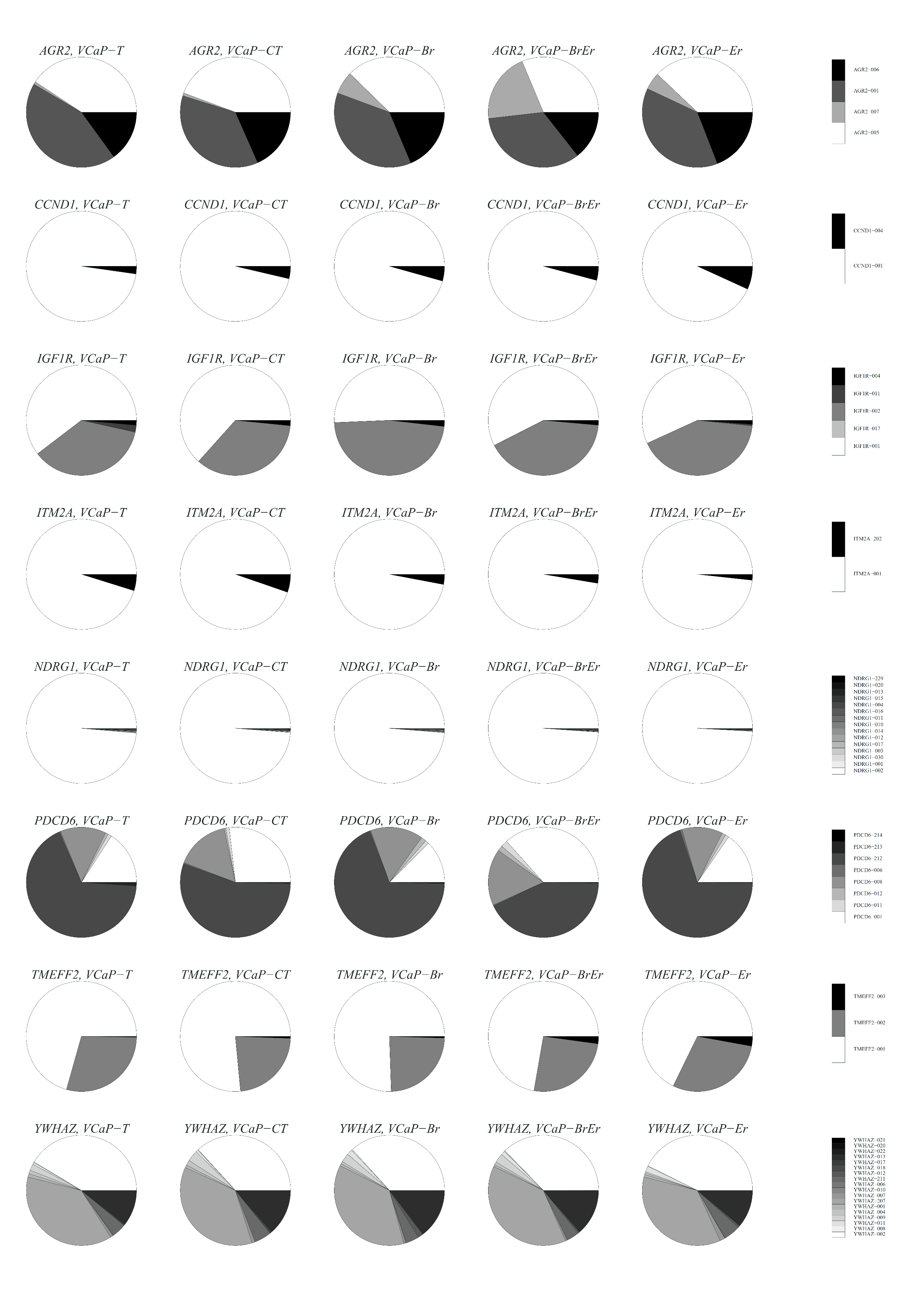

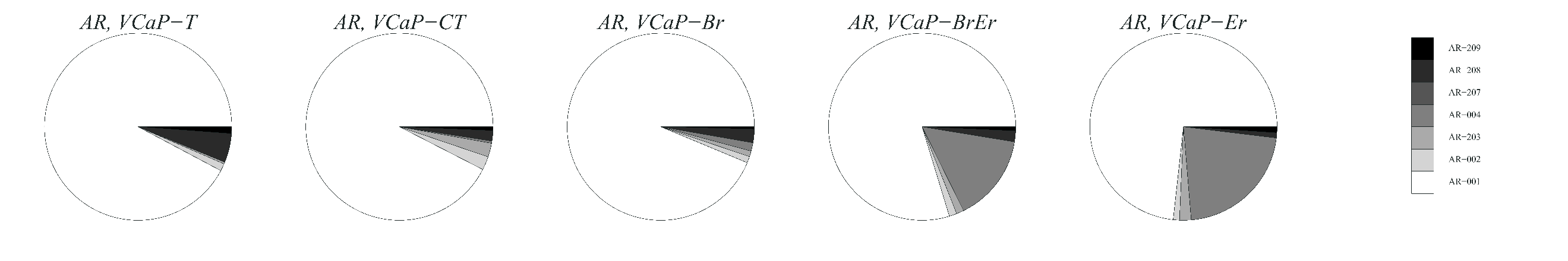

Supplementary figure 5. Transcriptomic splice variant profiles for *AGR2*, *CCND1*, *IGF1R*, *ITM2A*, *PDCD6,* *TMEFF2, YWHAZ*, *NDRG1* and *AR* genes. *AGR2* and *PDCD6* splice variant profiles show clear changes in VCaP-BR-ER cells whereas other cells are indifferent in terms of alternative splicing. *CCND1, IGF1R, ITM2A, TMEFF2, YWHAZ, and NDRG1* did not show major changes in their splice variant profile between any cell line. AR splice variant profile changes, by emerging of the AR-004, after the introduction of AR signalling inhibitor enzalutamide in both enzalutamide-resistant cell lines VCaP-BRER and VCaP-ER. Bicalutamide does not induce alternative splicing.
